# FER-mediated tyrosine phosphorylation and PIK3R2/p85β recruitment on IRS4 promotes the PI3K-AKT signaling pathway and tumorigenesis in ovarian cancer

**DOI:** 10.1101/2020.07.20.211417

**Authors:** Yanchun Zhang, Xuexue Xiong, Qi Zhu, Jiali Zhang, Yuetong Wang, Jian Cao, Li Chen, Shengmiao Chen, Linjun Hou, Xi Zhao, Piliang Hao, Min Zhuang, Dake Li, Gaofeng Fan

**Author notes:** Equal Contribution (Yanchun Zhang, Xuexue Xiong).

## Abstract

Tyrosine phosphorylation, orchestrated by tyrosine kinases and phosphatases, modulates a multi-layered signaling network in a time and space dependent manner. Dysregulation of this post-translational modification is inevitably associated with pathological diseases. Our previous work has demonstrated that non-receptor tyrosine kinase FER is upregulated in ovarian cancer. Knockdown of the kinase attenuates metastatic phenotypes in tumor cells. Here we employed mass spectrometry and biochemical approaches to identify IRS4 as a novel substrate of FER. Using a proximity-based tagging system, we determined that FER-mediated phosphorylation of Tyr779 enables IRS4 to recruit PIK3R2/p85β, the regulatory subunit of PI-3K, and activate the PI3K-AKT pathway. Rescuing IRS4-null ovarian tumor cells with phosphorylation-defective mutant, but not WT IRS4, delayed tumor cell proliferation both *in vitro* and *in vivo*. Overall, we revealed a kinase-substrate regulatory mode between FER and IRS4, and the pharmacological inhibition of FER kinase may be beneficial for ovarian cancer patients with PI3K-AKT hyperactivation.

## Introduction

Ovarian cancer is the most devastating gynecological malignancy, with high morbidity and ranking fifth among all cancer-related mortality in women (1). Patients with ovarian carcinoma are usually diagnosed at an advanced stage, resulting in a very low five-year survival rate (1). Despite recent advances in surgery, chemotherapy and radiotherapy for ovarian cancer patients, major medical challenges such as metastasis and therapeutic resistance remain largely unsolved. The lack of a wide-spread genetically-engineered mouse model for ovarian cancer significantly delays the entire process of ovarian cancer research. In particular, the molecular mechanisms of ovarian tumor progression and metastasis are not well understood, key factors severely restricting the overall survival from ovarian cancer (2). Consequently, there is an urgent need to and enormous translational potential in revealing the molecular mechanisms regulating the initiation, progression and metastasis of ovarian cancers. Understanding these mechanisms will serve as the first step toward identifying novel therapeutic targets and biomarkers for intervention against this heterogeneous and deadly disease (3).

Protein tyrosine kinases represent a family of important enzymes for controlling cell proliferation, motility, survival and differentiation, whose dysfunction have been closely related to the etiology of many major diseases, including cancer. Accordingly, tyrosine kinases are prominent drug targets. Depending on the cellular localization, the family can be further divided into receptor tyrosine kinase, which resides in the plasma membrane, and non-receptor tyrosine kinase, which resides in cytosol. Non-receptor tyrosine kinases are key regulators of intracellular signal transduction (4). The feline sarcoma kinase FES and feline sarcoma-related kinase FER represent a unique family of non-receptor tyrosine kinases. They are characterized by distinguishable N-terminal phospholipid-binding and a membrane targeting FER/CIP4 homology/Bin1/Amphiphysin/RVS (F-BAR) domain, which are reported to function in cell proliferation, motility, cell-to-cell adhesion, and mediate signal transmission from cell surfaces to the cytoskeleton (5, 6). Notably, the FER protein has been shown to be aberrantly upregulated and activated in different types of carcinoma (7–10). Specifically, high activity of FER protein kinase has been observed in 22% of ovarian cancer tumor samples via a global phosphoproteomic approach (10). To date, several receptor tyrosine kinases have been reported to act upstream of FER, including EGFR, PDGFR and integrin (6, 11, 12). Meanwhile, STAT3, cortactin and CRMP2, the functions of which are intensively involved in tumor cell motility and chemo-resistance, have been verified as FER substrates (13–15).

Previous studies in our laboratory have elucidated a dual mechanism for the regulation of MET through non-receptor protein tyrosine kinase FER in ovarian cancer (16, 17). The protein tyrosine kinase MET is the receptor of hepatocyte growth factor (HGF), distributes across cell membranes and is a target the treatment of ovarian carcinoma by molecular targeted therapy (18). MET overexpression and hyperactivation have been identified as a major contributing factor of ovarian tumorigenesis, malignant development, metastasis and overall poor prognosis (19). We have demonstrated that FER is significantly upregulated in ovarian cancer cell lines and ovarian cancer tumor samples, compared to normal controls and that its downregulation by RNAi results in substantial attenuation of tumor cell migration, invasion and metastasis (16). Mechanistically, we found that in the absence of ligand HGF, FER can directly phosphorylate the C-terminal recruitment region of MET and its key downstream scaffold protein GAB1, specifically activating SHP2–MAPK signaling and potentiating the migration and invasion phenotypes of ovarian cancer cells (16). Furthermore, in the presence of HGF, FER can sustain the distribution of MET on plasma membranes, thereby delaying its intracellular trafficking and inactivation by protein-tyrosine phosphatase PTP1B and enhancing kinase function of the receptor (17).

In addition to motility, FER expression has also been associated with proliferation of tumor cells (8, 20–23). However, due to the limited number of known substrates, the molecular basis for its pro-proliferation activity remains enigmatic. In this report, we aim to identify novel substrate(s) of FER and investigate the role of kinase-substrate regulatory modules in promoting ovarian tumorigenesis and progression. By integrating mass spectrometry analysis with biochemical and biological approaches, we have demonstrated that FER directly phosphorylates insulin receptor substrate 4 (IRS4) and that this tyrosine phosphorylation is important to create a binding site for recruiting PIK3R2/p85β. As the key regulatory subunit of PI-3K kinase, recruitment of PIK3R2/p85β onto IRS4 is required for PI3K-AKT signaling pathway activation and tumorigenesis in ovarian cancer.

## Results

### Mass spectrometry analysis identified IRS4 as a novel substrate of FER

In order to identify novel substrates for tyrosine kinase FER, we developed an overexpressed wild-type (FER-WT) and a kinase-dead mutant (FER-K592R) of FER, respectively, in human embryonic kidney 293FT (HEK293FT) cells. Tyrosine phosphorylated proteins in cells upon transfection were enriched by the anti-pTyr antibody, 4G10. We first separated these proteins by gel electrophoresis, followed by immunoblotting with anti-pTyr antibody (Fig 1A). We observed that significant numbers of proteins could be modified by tyrosine phosphorylation in cells expressing FER-WT compared to FER-K592R (Fig 1A). We excised the gel according to the position shown in Fig 1A and prepared these samples via in-gel tryptic digestion for mass spectrometry analysis. We obtained 2,298 candidate substrates with scores above 100 (Fig 1B and Supplemental Table S1.1~S1.6). Concurrently, immunoprecipitated proteins subjected to pTyr (4G10) antibody pull-down were directly digested on beads with trypsin and subjected to mass spectrometry. With this procedure we identified 153 candidate proteins (Fig 1B and Supplemental Table S1.7). Combining results from the two experiments we obtained 99 overlapped hits for further examination (Fig 1B).

**Figure 1.**
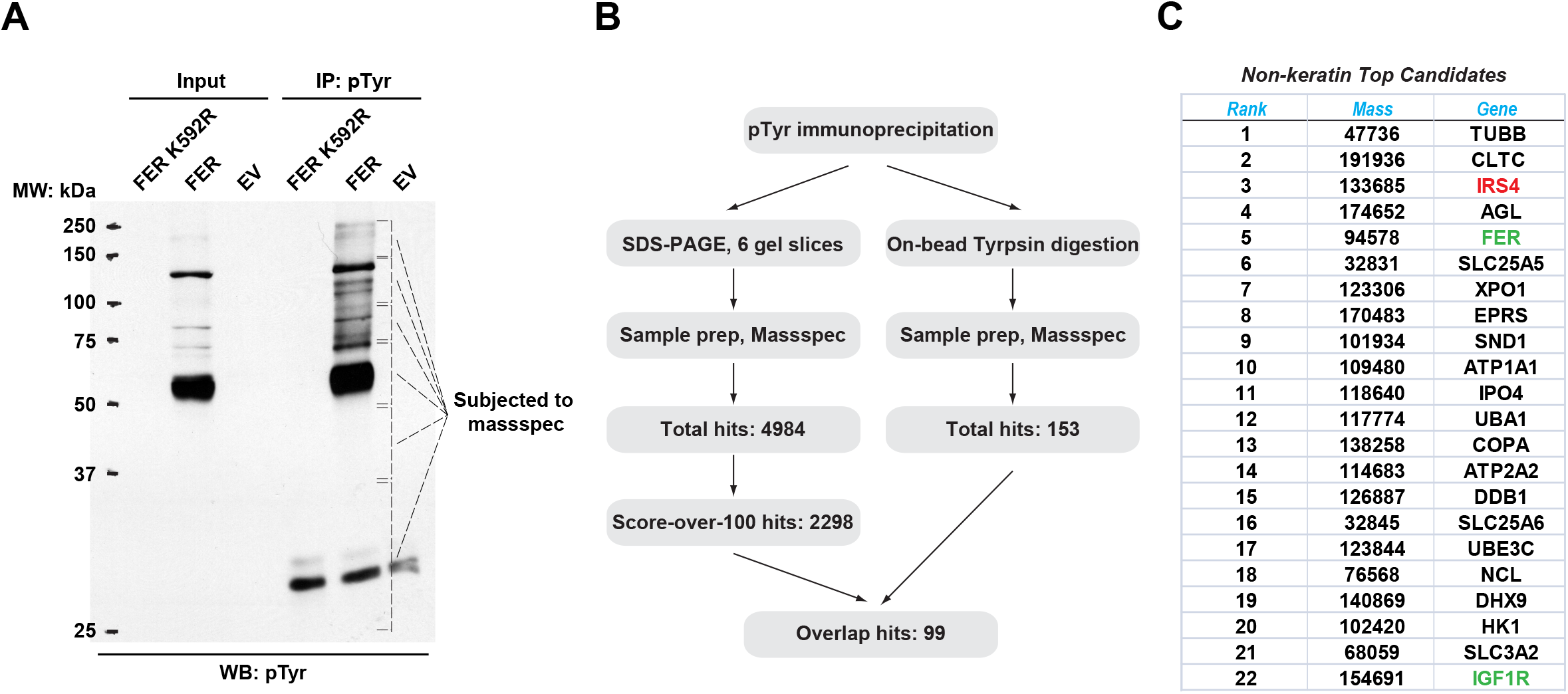
Mass spectrometry analysis identified IRS4 as a novel substrate of FER. A. FER-WT or FER-K592R was transfected into HEK293FT cells, followed by immunoprecipitation with antibody against pTyr. Immunopreciptates and whole cell lysates were subjected to immublotting analysis with anti-pTyr antibody. B. Experimental flow chart and results of two methods for potential substrates identification of FER kinase. The immunoprecipitated samples were subjected to in-gel (left) or on-bead (right) tryptic digestion, followed by mass spectrometry protein identification. C. Top-ranked overlapping candidate genes from both mass spectrometry analyses listed in B. Keratin genes were not shown.

The top 22 candidate genes are illustrated in Fig 1C and several reported FER substrates were included in the list. Notably, we identified FER as its own substrate since the Tyr402 residue of the kinase is known to be auto-phosphorylated (16). We also identified IGF-1R, a key tyrosine kinase receptor in regulating cancer cell survival, proliferation, and motility (24). In addition, a number of unreported genes were also listed, including XPO1 and IPO4 (cytoplasm-nucleus shuttle (25, 26)), SLC25A5, SLC25A6 and SLC3A2 (ADP/ATP transportation from mitochondria to cytoplasm, as well as heteromeric amino acid and polyamine transportation (27, 28)), and UBA1 and UBE3C (ubiquitin-proteasome degradation (29, 30)). In this study, we established gene IRS4 as our top candidate for the following reasons: (1) It was #3 in mass spectrometry score ranking. (2) A signaling molecule in the same pathway (IGF-1R) has been previously identified as substrate for FER kinase. (3) Tyrosine phosphorylation is an important post-translational modification in regulating the biological function of IRS4.

### FER kinase physically interacts with IRS4, involving the C-terminal region of FER and N-terminal region of IRS4

We first investigated if there was any physical interaction between FER and IRS4. We transiently expressed Myc-tagged IRS4 alone or together with FER in HEK293FT cells, followed by immunoprecipitation with resin against a Myc-tag. As shown in Fig 2A-B, IRS4 binds to FER and this association was not affected when FER-WT was replaced with its kinase dead mutant FER-K592R. Interestingly, the binding of GRB2 to IRS4 was FER kinase activity dependent (Fig 2B).

**Figure 2.**
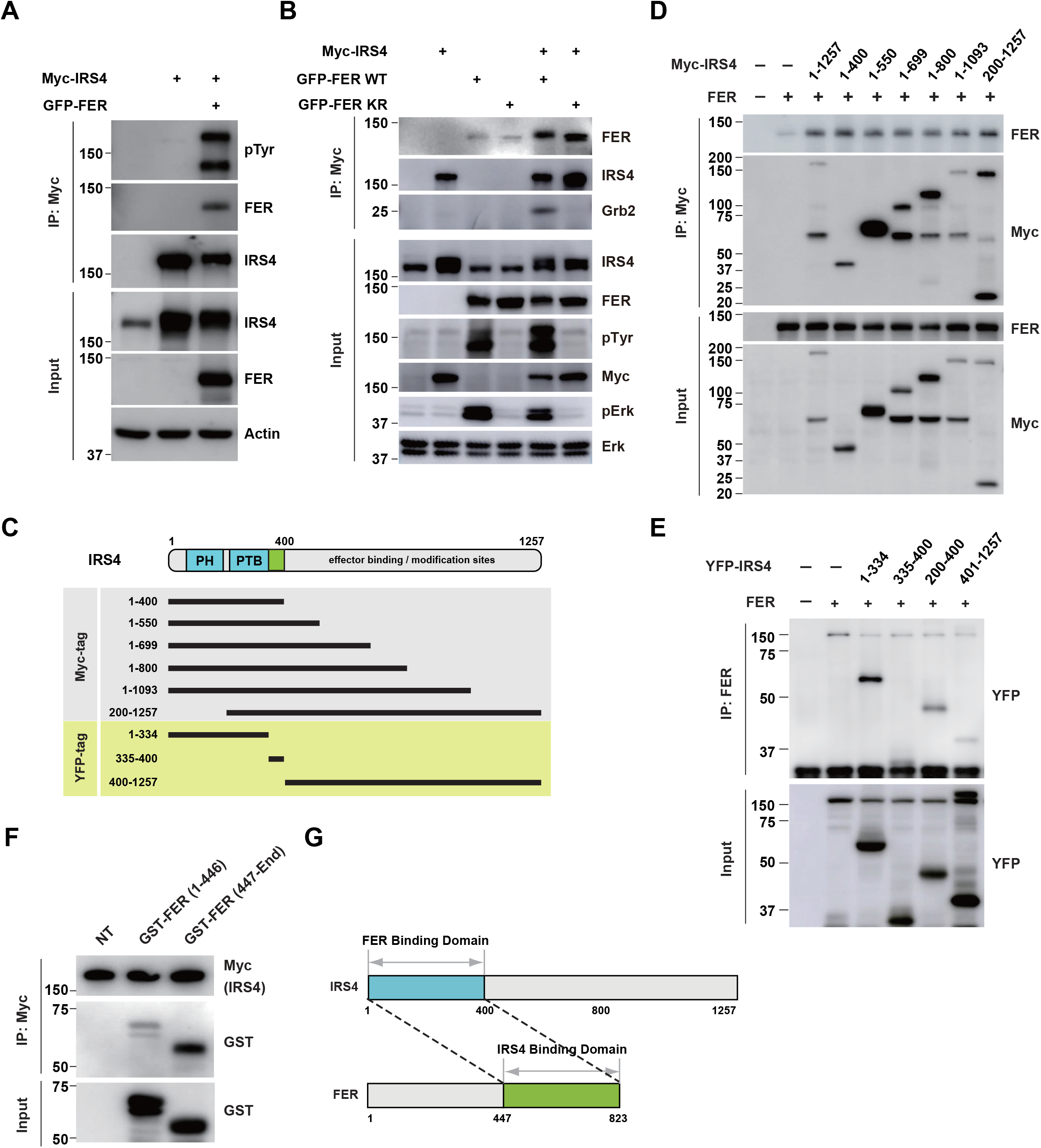
FER kinase physically interacts with IRS4, involving the C-terminal region of FER and N-terminal region of IRS4. A. Myc-tagged IRS4 were transiently transfected alone, or together with FER in HEK293FT cells, followed by immunoprecipitation with resin against Myc-tag, and immunobloting with IRS4, FER, 4G10 antibodies, respectively. The interaction of FER with IRS4 was examined. Expressions of IRS4, FER and loading control actin in whole cell lysate samples were also probed. B. After transient transfection with Myc-tagged IRS4, FER-WT or KR mutant in HEK293FT cells, IRS4 was immunoprecipitated with resin against Myc-tag from cell lysates and probed for association with FER and GRB2. The blot was also probed with antibodies against IRS4, FER, 4G10, Myc, phosphor- and total-ERK in whole cell lysate samples. C. Schematic illustration of domain structure of IRS4. PH: pleckstrin homology domain; PTB: phosphotyrosine binding domain. Multiple truncation mutants of IRS4, including fragments corresponding to amino-acid residues 1-400, 1-550, 1-699, 1-800, 1-1093, 200-1257, 1-334, 335-400, 200-400 and 401-1257, with Myc or YFP tag were also listed. D. FER was expressed alone or co-expressed with a series of Myc-tagged truncated forms of IRS4 in HEK293FT cells, as indicated. Lysates were harvested and immunoprecipitated with anti-Myc resin. The associations of FER with different IRS4 truncation mutants were compared. Expressions of IRS4 and FER in whole cell lysate samples were also probed. E. HEK293FT cells were transiently co-transfected with GFP-tagged FER and truncated versions of IRS4 with YFP tag, followed by immunoprecipitation with FER antibody and immunobloting with YFP antibody. F. Two truncated FER protein with GST tag were expressed and purified from E.coli. GSH-beads bound recombinant proteins were further incubated with whole HEK293FT cell lysates expressing Myc-tagged IRS4, respectively. Immunoprecipitates were subjected to immunoblotting analysis to assess the region requirement for IRS4 interaction. G. Schematic illustration of regions involved in binding between IRS4 and FER.

To map the particular region(s) involved in the interaction between IRS4 and FER, we employed different truncation forms of IRS4 with a Myc-tag, including fragments corresponding to amino-acid residues 1-400, 1-550, 1-699, 1-800, 1-1093 and 200-1257, as indicated in Fig 2C. These truncation mutants were overexpressed when combined with FER in HEK293FT cells and the binding between them and FER was detected by immunoprecipitation (Fig 2D). Surprisingly, the shortest truncation of IRS4 (1-400) maintained a similar binding affinity with FER, indicating the N-terminal region of IRS4 is crucial for the interaction (Fig 2D).

To further pinpoint the key region involved in binding, we employed additional truncated versions of IRS4 with a YFP-tag, including segments with amino-acid residues 1-334, 335-400, 200-400 and 401-1257 (Fig 2C) and assessed the binding of FER to different IRS4 fragments by immunoprecipitation (Fig 2E). Whereas both the 335-400 and 401-1257 mutants had very limited binding affinity with FER, the N-terminal mutant 1-334 showed strongest binding among all these truncated constructs (Fig 2E). Additionally, the 200-400 truncated mutant of IRS4 also illustrated weak but significant binding affinity with FER (Fig 2E). These results highly suggested that the 1-334 residues of IRS4 are essential for its association with the kinase FER.

Subsequently, we expressed and purified two truncated FER proteins with a GST-tag in *E.coli* (Fig 2F) and incubated them separately with whole HEK293FT cell lysates expressing Myc-tagged IRS4 to evaluate their interaction. The C-terminal region (447-823) of FER, rather than its N-terminal region (1-446), showed robust interaction with IRS4 (Fig 2F). This result demonstrates that FER physically interacted with IRS4, and taken with the previous result shows that C-terminal region of FER and N-terminal region of IRS4 are critical for binding (Fig 2G).

### Mass spectrometry analysis and site-directed mutagenesis identified several FER-phosphorylated tyrosine residues on IRS4

We next investigated the molecular details in the tyrosine phosphorylation of IRS4 by FER kinase. The first question we wanted to address was the specificity of regulation. To this end, we overexpressed 7 non-receptor tyrosine kinases in parallel and assessed the phosphorylation extent change of IRS4 post immunoprecipitation followed by anti-phosphotyrosine blotting analysis. As shown in Fig 3A, among all these kinases, FER illustrated highest capability for tyrosine phosphorylation of IRS4.

**Figure 3.**
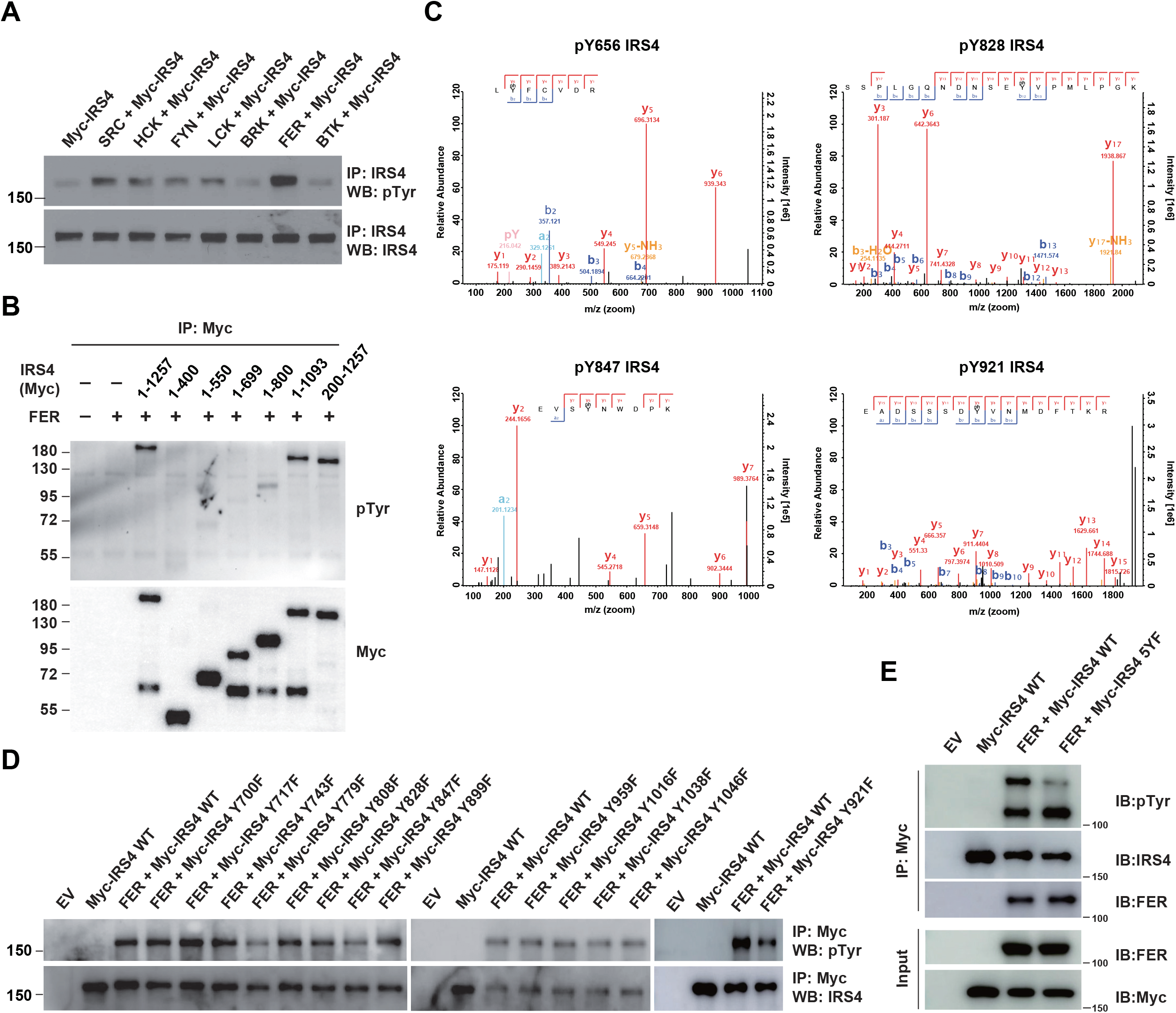
Mass spectrometry analysis and site-directed mutagenesis identified several FER-phosphrylated tyrosine residues on IRS4. A. Myc-IRS4 was transfected alone, or together with several protein-tyrosine kinases in HEK293FT cells, as indicated. Myc-IRS4 was immunoprecipitated with resin against Myc-tag and the phosphorylation level of IRS4 was examined by pTyr (4G10) immunobloting. B. After transient transfection of a series of Myc-tagged different truncated forms of IRS4 along with FER in HEK293FT cells, as indicated, lysates were harvested and immunoprecipitated with anti-Myc resin. The tyrosine phosphorylation level of IRS4 truncations were examined with antibody aginst pTyr (4G10). C. Results of mass spectrometry analysis identified 4 tyrosine residues (Y656, Y828, Y847 and Y921) of IRS4 phosphorylated by FER kinase. D-E. HEK293FT cells were transiently transfected with Myc-tagged IRS4 or its YF mutants with FER, as indicated, followed by immunoprecipitation with resin against Myc-tag and immunobloting with IRS4, FER, pTyr (4G10) antibody. The phosphorylation levels of different IRS4 mutants were compared to illustrate the effects of point mutations in IRS4 on its phosphorylation by FER

To determine which region on IRS4 can be phosphorylated by FER, we overexpressed abovementioned Myc-tagged IRS4 truncation mutants in conjunction with FER in HEK293FT cells (Fig 3B) and evaluated the phosphorylation extent change of IRS4 fragments. We concluded tyrosine residues between 700 and 1093 amino acids of IRS4 were potential substrate(s) for FER kinase, since: (1) the 1-1093 mutant of IRS4 showed equivalent phosphorylation level to full-length IRS4 upon FER phosphorylation; (2) compared to the 1-699 mutant, 1-800 mutant of IRS4 showed weak but significant amount of phosphorylation.

We took two strategies to further pinpoint the particular tyrosine residue(s) that undergo phosphorylation in the presence of FER. By performing mass spectrometry analysis in duplicate, we observed 3 tyrosine sites, namely Y656, Y828 and Y921, whose phosphorylation were repeatedly detected in both datasets (Fig 3C and Supplemental Table S2.1~2.2). Y847 was also detected once (Fig 3C and Supplemental Table S2.1). Interestingly, these four residues are within or very close to the 700~1093 region of IRS4. Afterwards, we mutated all tyrosine residues within 700~1093 region of IRS4, including Y700, Y717, Y743, Y779, Y808, Y828, Y847, Y899, Y959, Y1038 and Y1046, to phenylalanine and assessed the phosphorylation level change of these mutants compared to WT IRS4. Excluding the Y828, Y847 and Y921 mutants, a tyrosine to phenylalanine substitution at residue 779 remarkably decreased the phosphorylation level of IRS4 in the presence of FER (Fig 3D).

By combining results from both mass spectrometry analysis and site-directed mutagenesis analysis, we generated a quintuple Tyr to Phe mutant of IRS4, including Y656, Y779, Y828, Y847 and Y921 (Named “5YF” mutant hereafter). As is consistent with our previous observations, the tyrosine phosphorylation level of 5YF mutant by FER was profoundly diminished compared to each single mutant (Fig 3E). These results highly suggest that there are five major tyrosine residues within IRS4 that are subjected to FER-mediated phosphorylation.

### IRS4 was upregulated in certain ovarian carcinoma-derived cell lines and the loss of IRS4 inhibited PI3K-AKT pathway activation and ovarian cancer cell proliferation

Previous studies from our lab have demonstrated the aberrantly high expression of FER kinase in ovarian cancer and its important role in promoting tumor cell metastasis both *in vitro* and *in vivo* (16). In this study we want to adapted the same cell model to further evaluate the biological function of the FER-IRS4 kinase-substrate pair in a physiological context. We first used immunoblotting analysis to compare the expression level of the IRS4 protein between 2 human ovarian surface epithelial (HOSE) cell lines immortalized by the human papilloma viral oncogenes E6 and E7, and 11 ovarian carcinoma-derived cell lines. Compared to normal HOSE control cells, two ovarian carcinoma-derived cell lines, OVCAR-5 and OVCAR-3, showed evident up-regulation of IRS4 protein expression (Fig 4A). Of note, FER is also upregulated in both cell lines (16).

**Figure 4.**
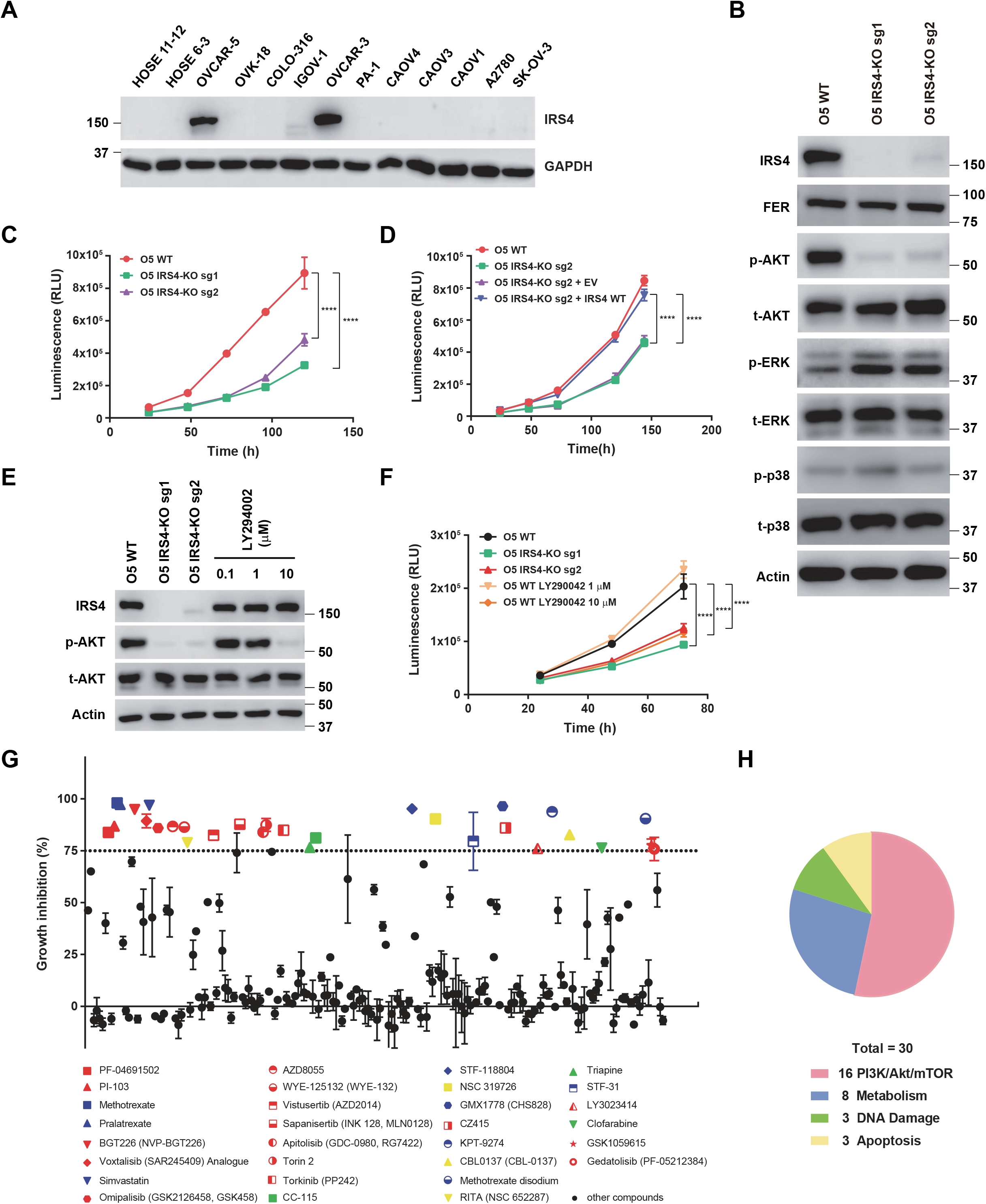
IRS4 was upregulated in certain ovarian carcinoma-derived cell lines and the loss of IRS4 inhibited PI3K-AKT pathway activation and ovarian cancer cell proliferation. A. Immunobloting of IRS4 with two immortalized human ovarian surface epithelial (HOSE) cell lines and 11 ovarian carcinoma-derived cell lines to demonstrate increased expression level of IRS4 in certain ovarian carcinoma-derived cell lines. GAPDH was probed as the loading control. B. Immunoblotting analysis to illustrate the knockout effect of IRS4 on activation of downstream signaling pathways. Sample lysates were prepared from parental and IRS4-KO OVCAR-5 cell lines, and immunoblotted with the designated antibodies, including IRS4, pS473- and total-AKT, phosphor- and total-ERK, and phosphor- and total-p38 antibodies, with actin as loading control. C. CellTiter-Glo luminescent cell viability assay with OVCAR-5 WT cells and OVCAR-5 IRS4-KO cells at the indicated time intervals. Results represent mean ± S.D. ****P < 0.0001. D. CellTiter-Glo cell viability assay was conducted to evaluate cell proliferation after restoration of IRS4 expression in OVCAR-5 IRS4-KO cells. The ectopic expression of empty vector (EV) was used as a negative control. Results represent mean ± S.D. ****P < 0.0001. E. Lysates from OVCAR-5 WT and IRS4-KO cells were harvested and the expression of phospho- and total AKT was detected by immunobloting. AKT inhibitor LY294002 (0.1 μM, 1 μM, and 10 μM), as indicated, were incubated for 24 hours. F. Cell proliferation was assessed in OVCAR-5 cells following IRS4 knockout (IRS4-KO sg1 or sg2) or incubation with AKT inhibitor LY294002 (1 μM and 10 μM) using the CellTiter-Glo luminescent cell viability assay at the indicated time intervals. Results represent mean ± S.D. ****P < 0.0001. G. CellTiter-Glo cell viability assay was used to assess proliferation inhibition in OVCAR-5 cells following the treatment of 198 metabolic targeting compounds (1 μM) in compound library from Selleck. The ratios of luminescence in treated OVCAR-5 cells to that in untreated OVCAR-5 cells were calculated as the percentage of proliferation inhibition. Dashed lines indicate 75% proliferation inhibition percentage. All compounds above this threshold are represented as color dots: red dots represent compounds targeting PI3K/AKT/mTOR pathway, blue dots represent compounds targeting metabolism-related pathway, green dots represent compounds targeting DNA damage-related pathway and yellow dots represent compounds targeting apoptosis-related pathway. Data are shown as means ± SEM. H. Pie chart summary representing the proportion of different signaling pathway targeted by the compounds with over 75% proliferation inhibition percentage (%) in OVCAR-5 cells, where red color represent 16 compounds targeting PI3K/AKT/mTOR pathway, blue color represent 8 compounds targeting metabolism-related pathway, green color represent 3 compounds targeting DNA damage-related pathway and yellow color represent 3 compounds targeting apoptosis-related pathway.

Next, we applied the CRISPR-Cas9 system to genetically knockout the IRS4 gene in ovarian cancer cells and investigated the change of cellular phenotype(s) after IRS4 deletion. The knockout effect of IRS4 was confirmed via immunoblotting against an anti-IRS4 antibody. Regrettably, after several attempts we were unable to get a single clone of IRS4 knockout OVCAR-3 cells, since this particular cell line prefers to grow in clusters and it cannot re-confluence after FACS sorting procedure. However, we successfully obtained two OVCAR-5 IRS4 knockout cell lines with two distinct sgRNAs, (Fig 4B).

We then assessed the changes in cancer cell fitness in the absence of IRS4. Interestingly, Cell Titer-Glo (CTG) luminescent cell viability assay demonstrated that the knockout of IRS4 notably delayed the proliferation capacity of OVCAR-5 cells (Fig 4C). Conversely, ectopic re-expression of IRS4, rather than an empty vector, restored the potential of cell proliferation in IRS4-KO ovarian cancer cells (Fig 4D), highlighting the key role of IRS4 in maintaining ovarian tumor cell growth.

We explored further and compared the difference(s) in downstream signaling pathways upon IRS4 deletion. Whereas there was little change in phospho-Erk and phospho-p38 levels, we observed a dramatic decrease of phospho-AKT in OVCAR-5 IRS4-KO cells (Fig 4B), indicating a tight correlation between PI3K-AKT pathway and OVCAR-5 cancer cell proliferation. To confirm this, we treated OVCAR-5 WT cells with AKT inhibitor LY294002 (Fig 4E-F) and demonstrated that the inhibition of the PI3K-AKT pathway in OVCAR-5 cells did indeed decrease the phosphorylation level of AKT and cell growth in a concentration-dependent manner. These results revealed an important role of IRS4 in regulating the PI3K-AKT signaling pathway and cell proliferation of OVCAR-5 ovarian cancer cells.

To further address the importance of PI3K-AKT signaling pathway in regulating ovarian cancer cell growth, we performed a CTG-based compound library screening assay in OVCAR-5 WT ovarian cancer cells. This compound library contains 198 small molecule inhibitors targeting key node genes in cancer cell signaling transduction and metabolism. Briefly, OVCAR-5 cells were treated with each compound at concentration of 1 μM; three days later, CTG assay was applied to measure the cells proliferation rate. 30 out 198 compounds showed proliferation inhibition greater than 75% (Fig 4G-H). Among these, 16 compounds targeted the PI3K-AKT-mTOR signaling pathway while the remaining compounds targeted a variety of other pathways, including metabolism, DNA damage response and apoptosis. This result further supports the supposition that the PI3K-AKT-mTOR signaling pathway plays a critical role in controlling the proliferation of OVCAR-5 ovarian cancer cells, and it is of great interest to understand the molecular details of how IRS4 mediates AKT activation.

### PIK3R2 was identified as one of the major downstream signaling components of IRS4 by proximity labeling PUP-IT assay

Encouraged by our previous results we attempted to delineate the molecular mechanism by which IRS4 regulates the AKT signaling pathway, and therefore the likely mechanism by which it regulates the cell proliferation of ovarian cancer. We postulated that FER-mediated IRS4 phosphorylation would facilitate signaling molecule recruitment, which is important for AKT activation. To assess this hypothesis, we applied Pupylation-based interaction tagging (PUP-IT) system to identify potential interacting proteins of phosphorylated IRS4. PUP-IT has been reported as a new ligase-mediated proximity labelling technique, with advantages in detecting weak and transient protein-protein interactions under physiological conditions in living cells (31). Briefly, IRS4-WT or IRS4-YF mutant genes were fused to the PafA ligase which can covalently attach a small protein tag biotin-Pup (E) to substrate lysine residues of IRS4 binding proteins. All biotin-Pup (E) labeled candidate proteins were enriched with streptavidin beads and identified by mass spectrometry analysis (Fig 5A). Specifically, we focused on the differences in binding proteins between IRS4-WT and multiple IRS4-YF in OVCAR-5 cells. In combination with results from previous mass spectrometry and site-directed mutagenesis assays, we included five single Y to F mutants (Y779F, Y847F, Y921F, Y656F and Y828F) into this PUP-IT study. We also included the IRS4-5YF mutant, in which all five abovementioned Tyr residues were replaced by Phe.

**Figure 5.**
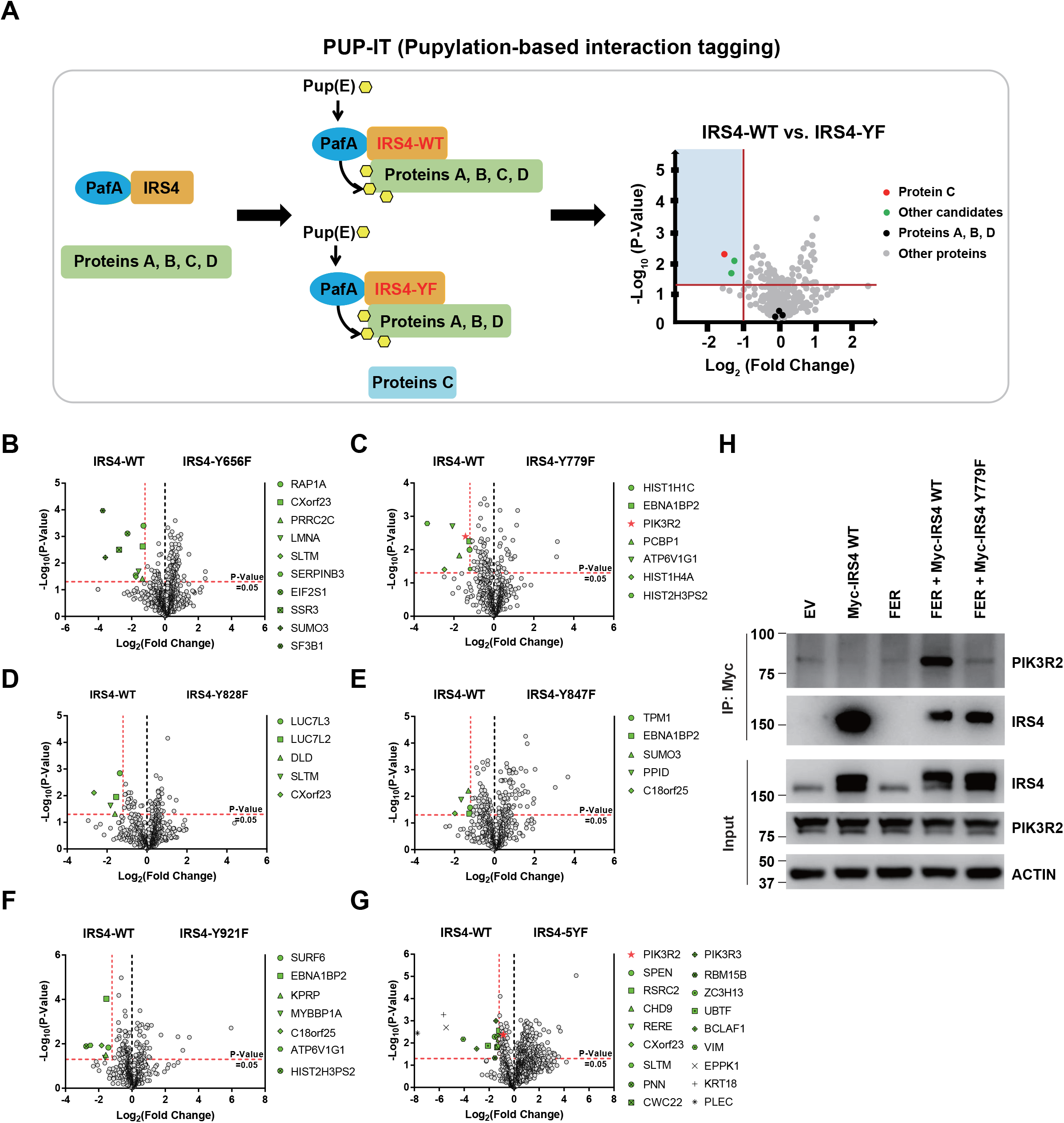
PIK3R2 was identified as one of the major downstream signaling components of IRS4 by proximity labeling PUP-IT assay. A. Scheme of potential interacting proteins identification for tyrosine phosphorylated IRS4 using PUP-IT proximity labeling platform. Final result is illustrated with volcano plots drawn by GraphPad Prism. B-G. PUP-IT results from mass spectrometry analysis for each potential interacting protein were showed in volcano plots, with LFQ intensity fold change (x-axis) and P-value change (y-axis). Fold change > 2.3 and P-value < 0.05 (T-test) were regarded as difference and statistical significance. Genes with significant binding differences were presented as green or red (PIK3R2) dots. H. Myc-tagged IRS4 or its Y779F mutant was co-expressed with FER in HEK293FT cells, as indicated, followed by immunoprecipitation with resin against Myc-tag and immunobloting with IRS4 and PIK3R2 antibodies. The blot was also probed with antibodies against IRS4, PIK3R2 and ACTIN in input samples.

The results from mass spectrometry analysis were summarized and illustrated as volcano map (Fig 5B-G), with Log_2_ (Fold Change) as the x-axis and −Log_10_ (P-value) as the y-axis. For each plot, we highlighted genes with P-value less than 0.05 (-Log_10_ (P-value) = 1.30) and preferential IRS4-WT binding over IRS4-YF mutant, with fold change greater than 2.30 (Log_2_ (Fold Change) = −1.20). Among all detected proteins, the number of unique peptides of IRS4 ranked at the top (Supplemental Table S3.1~S3.3), indicating the robustness of the assay. Compared to the IRS4-WT control, three YF mutants of IRS4 (Y779F, Y847F and Y921F) showed reduced binding affinity with EBNA1 Binding Protein 2 (EBNA1BP2) (Fig 5C and Fig 5E-F). SUMO3, SLTM, ATP6V1G1, HIST2H3PS2, CXorf23 and C18orf25 also showed reduced binding affinity with two of the YF mutants of IRS4. Notably, we observed that PIK3R2 (the regulatory subunit of PI-3K) preferred to bind with WT rather than the Y779F mutant of IRS4 (Fig 5C). Interestingly, this preferential association of PIK3R2 appeared again when comparing interaction proteins between WT and 5YF mutant of IRS4 (Fig 5G). The results from these PUP-IT assays strongly suggest that the FER-triggered tyrosine phosphorylation of IRS4 facilitated the recruitment of PIK3R2 and that Tyr779 was the major contributing residue.

To further validate the PUP-IT results, we performed an immunoprecipitation experiment to assess the essentiality of IRS4 phosphorylation on Y779 site in recruiting PIK3R2. FER was transfected, alone or together with Myc-IRS4 (WT) or Myc-IRS4 (Y779F), followed by anti-Myc immunoprecipitation and PIK3R2 detection (Fig 5H). Compared to the WT control, the Y779F mutation of IRS4 exhibited dramatically decreased binding affinity with PIK3R2 (Fig 5H), indicating the importance of the tyrosine phosphorylation of the 779 residue (as mediated by FER) in recruiting PIK3R2. PIK3R2 (also known as p85β) is a key regulatory subunit of PI3K in the PI3K-AKT signaling pathway (32). Unlike PIK3R1 (p85α) whose function is tumor suppressive, PIK3R2 plays a role as oncogene (32). Both PUP-IT assay and biochemical pulldown assay consistently demonstrated that FER-mediated tyrosine phosphorylation of IRS4 at Tyr779 enhanced the recruitment of PIK3R2 and activation of the PI3K-AKT signaling pathway, providing new insights into signaling events that underlie cell proliferation in ovarian carcinoma cells.

### FER-mediated PIK3R2 recruitment by IRS4 is crucial to ovarian cancer cell proliferation *in vitro* and tumorigenesis *in vivo*

To further pursue the role of the kinase FER in phosphorylating IRS4 in ovarian cancer cells, we generated FER knockout OVCAR-5 ovarian cancer cells by CRISPR-Cas9 and tested whether the global tyrosine phosphorylation of IRS4 would be affected upon FER loss. We observed decreased tyrosine phosphorylation of IRS4 after tandem IRS4 immunoprecipitation and pTyr immunoblotting (Figure 6A). Interestingly, the association of IRS4 and PIK3R2 was also decreased in FER-deficient OVCAR-5 ovarian cancer cells (Figure 6A).

**Figure 6.**
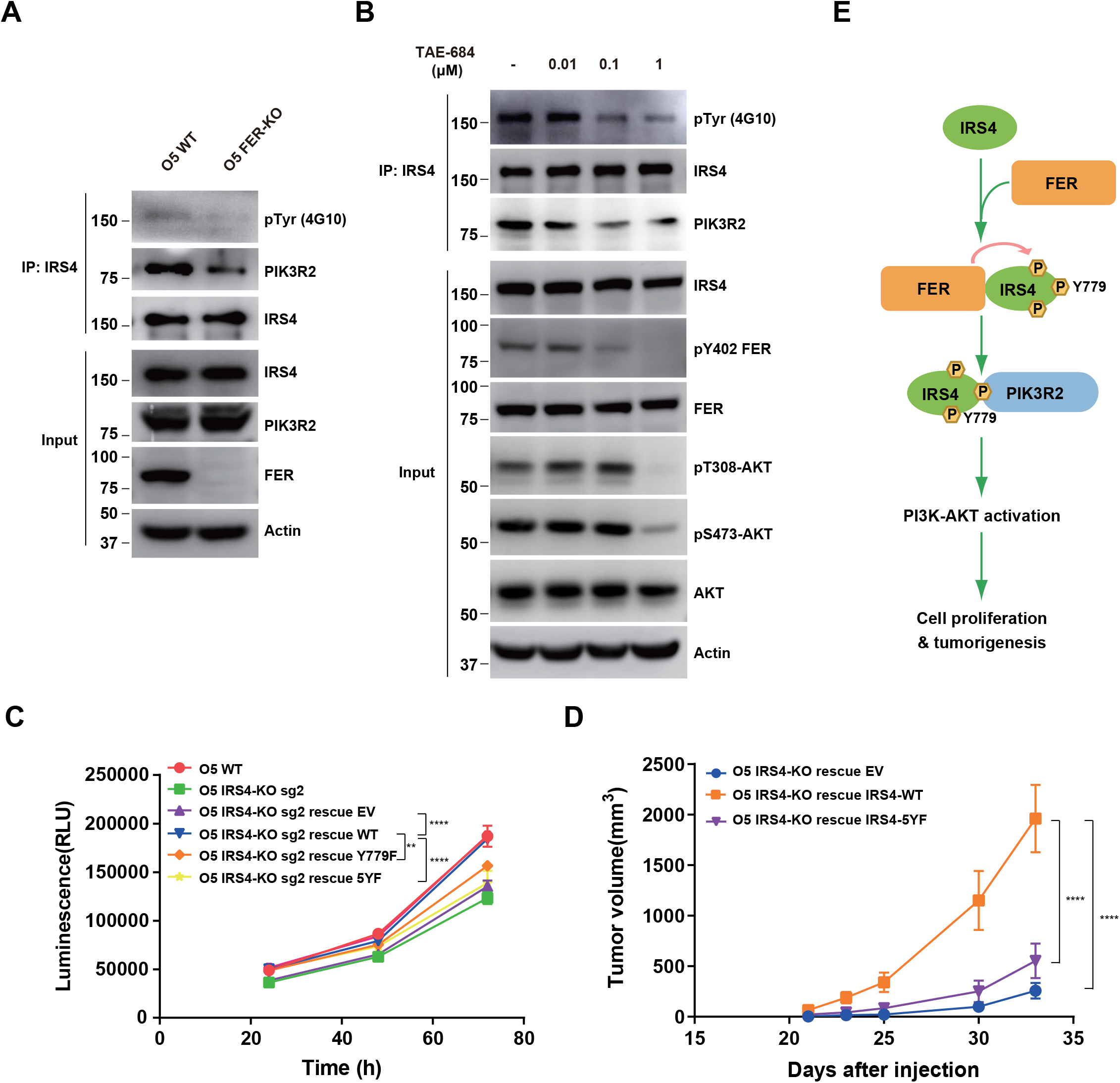
FER-mediated PIK3R2 recuritment by IRS4 is crucial to ovarian cancer cell proliferation *in vitro* and tumorgenesis *in vivo*. A. OVCAR-5 WT and OVCAR-5 FER-KO cell lysates were harvested and immunoblotted for pTyr (4G10), PIK3R2 and IRS4. Actin was probed as loading control. After IRS4 was immunoprecipitated from cell lysates, the global tyrosine phosphorylation of IRS4 and co-immunuoprecipitation of PIK3R2 were examined with pTyr (4G10) and PIK3R2 antibodies. B. OVCAR-5 cells treated with TAE684 (0.01 μM, 0.1 μM, and 1 μM) for 24 hours were lysed, and the expressions of IRS4, pY402 FER, FER, p-AKT and Actin were detected by immunobloting as indicated. IRS4 immunoprecipitation and pTyr immunoblotting was also used to test tyrosine phosphorylation level of IRS4 upon treatment. C. CellTiter-Glo cell viability assay was conducted to evaluate cell proliferation after rescue expression of WT or YF IRS4 in OVCAR-5 IRS4-KO cells. OVCAR-5 IRS4-KO cells, and OVCAR-5 IRS4-KO cells expressing empty vector (EV) were included as negative controls. The parental OVCAR-5 cells was used as a positive control. Results represent mean ± SEM. **P = 0.0024; ****P < 0.0001. D. After subcutaneous injection of OVCAR-5 IRS4-KO cells rescued with EV (n=5), IRS4-WT (n=4) and IRS4-5YF (n=7), respectively in the xenograft NSG mouse model, tumor volumes (in cubic millimeters, formula: volume=width2×length/2) were measured with calipers at the indicated time intervals. Results represent mean ± SEM. ****P < 0.0001. E. Working model: FER binds directly to IRS4, and phosphorylates its several tyrosine residues. FER-mediated phosphorylation of Tyr779 on IRS4 enhances recruitment of PIK3R2/p85β, the regulatory subunit of PI-3K, and promotes PI3K-AKT signaling pathway, which eventually leading to cell proliferation and tumorigenesis in ovarian cancer.

In a screen of 586 compounds, TAE684 has been identified as a potent inhibitor against FES, the family member of FER (33). The high similarity between FER and FES inspired us to evaluate if TAE684 exhibits an equivalent inhibitory effect on FER. Indeed, ovarian cells treated with TAE684 showed dose-dependent inhibition on the kinase activity of FER, as illustrated by pY402 FER blotting analysis and the activation of AKT (Fig. 6B). Meanwhile, results from tandem IRS4 immunoprecipitation and pTyr immunoblotting again indicate the reduced global tyrosine phosphorylation of IRS4 upon TAE684 treatment (Fig. 6B). Collectively, genetic ablation or pharmacological inhibition of tyrosine kinase FER in ovarian cancer cells leads to decreased global tyrosine phosphorylation and PIK3R2 recruitment of IRS4, which is consistent with our ectopic over-expression studies in HEK293FT cells (Figure 2A).

Our previous observations have already demonstrated a retarded cell proliferation in IRS4-deficient ovarian cancer cells (Fig 4C-D). To further evaluate the importance of tyrosine phosphorylation of IRS4, we re-expressed these Y to F mutants, along with the WT, in OVCAR-5 IRS4-KO ovarian cancer cells. Whereas re-expression of WT IRS4 in IRS4-KO cell line fully recovered the growth defect, 5YF mutants of IRS4 failed to rescue cell proliferation (Fig 6C). We did observe a certain extent of growth defect rescue with cells re-expressing in the Y779F mutant of IRS4, but these cells still showed significantly delayed proliferation rate compared to the parental ovarian cancer cells (Fig 6C, P = 0.0024). These results demonstrate that the FER kinase mediated tyrosine phosphorylation of IRS4 plays a key function in controlling cell proliferation in ovarian cancer.

The significant difference between the WT and YF mutants of IRS4 in regulating cell proliferation *in vitro* prompted us to extend the comparison of ovarian tumorigenesis *in vivo*. We adopted a xenograft mouse model with subcutaneous injection of OVCAR-5 IRS4-KO cells which was rescued with either an empty vector, or a WT or 5YF mutant of IRS4. Compared to mice injected with OVCAR-5 IRS4-KO cells rescued with WT IRS4, we observed significantly delayed tumor formation (P < 0.0001) in mice injected with OVCAR-5 IRS4-KO cells with an empty vector (Fig 6D). Consistent with our previous findings in cell cultures, the tumor growth in mice injected with OVCAR-5 IRS4-KO cells rescued with 5YF IRS4 were profoundly delayed compared to WT control (Fig 6D), further emphasizing the necessity of the tyrosine phosphorylation of key residues in IRS4 for ovarian tumorigenesis and progression.

### Aberrantly high expression of IRS4 was inversely correlated with prognosis in patients with ovarian cancer

To investigate the expression pattern of IRS4 in human organs and tissues, we first analyzed the RNA-seq data from Human Protein Atlas (HPA) dataset (http://proteinatlas.org). Interestingly, IRS4 shows the highest mRNA transcript abundance in the ovaries, followed by the thyroid gland and endometrium (Figure 7A). We further compared the protein expression levels of IRS4 among tissue microarrays of both normal ovaries and malignant ovarian carcinoma in the HPA database. In line with our findings in cell cultures, we observed a higher expression of IRS4 in ovarian cancer patient samples (Figure 7B).

**Figure 7.**
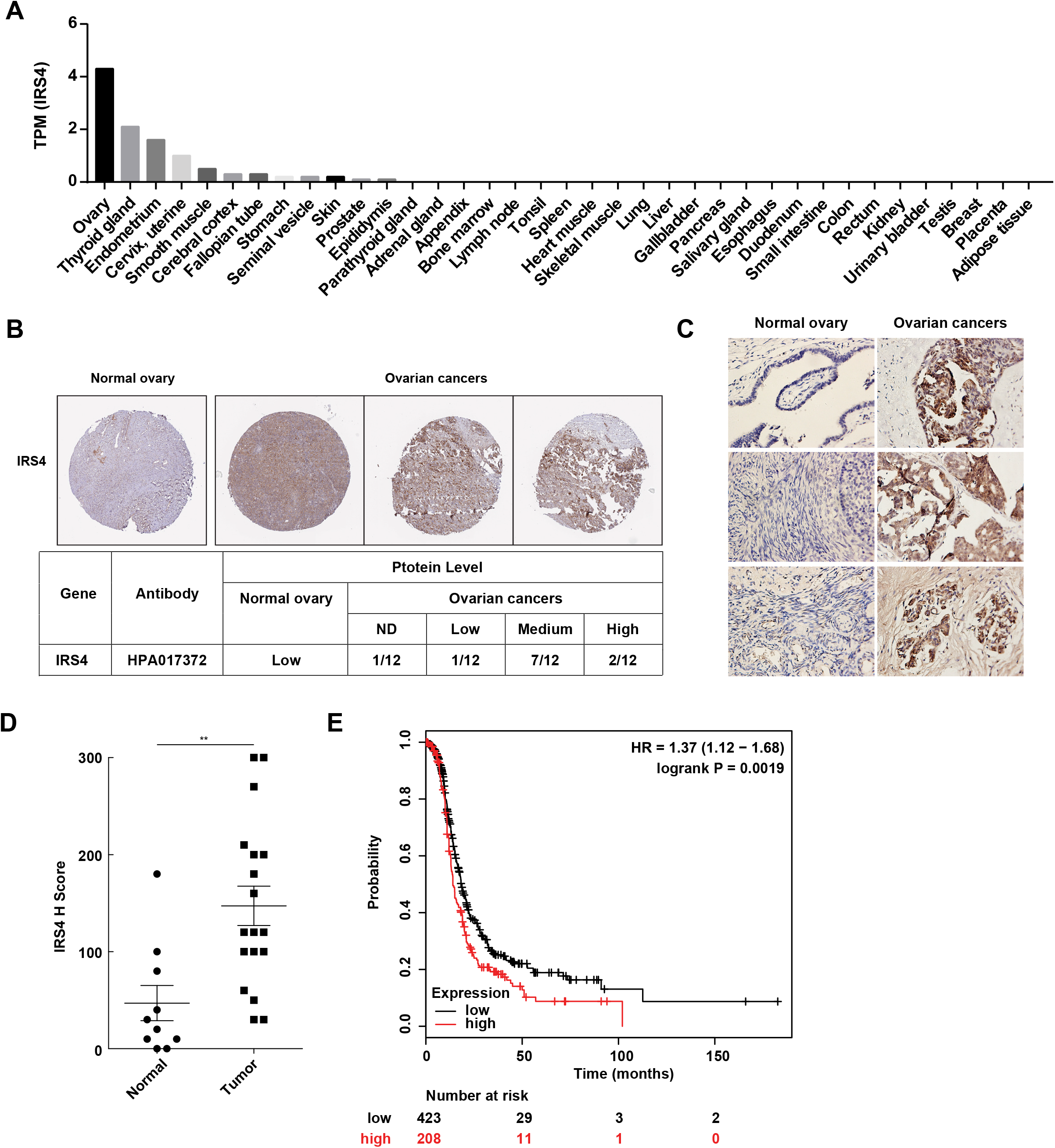
Aberrantly high expression of IRS4 was inversely correlated with prognosis in patients with ovarian cancer. A. mRNA expression profile of IRS4 in multiple human tissues based on RNA-seq tissue data from the HPA dataset (http://proteinatlas.org). Data is reported as mean pTPM (protein-coding transcripts per million), corresponding to mean values of the different individual samples from each tissue. B. The expression of IRS4 in the sections of normal ovary and malignant ovarian carcinoma according to representative tissue microarrays cores from the HPA database. C. Immunohistochemistry staining for IRS4 protein in normal ovaries (n=10) and malignant ovarian carcinomas (n=18) samples. Representative images are shown. D. Summary and statistical analysis of immunohistochemistry staining status of IRS4 H score between normal ovaries (n=10) and malignant ovarian carcinomas (n=18) samples. Results represent mean ± S.D. **P = 0.0029. E. Overall survival from over 600 ovarian cancer patients among previously published data sets by the km Plotter (http://www.kmplot.com). IRS4 expression is stratified as high versus low against median expression.

In addition, we collected 10 cases of normal ovary samples and 18 cases of malignant ovarian carcinomas samples to further explore the expression differences in IRS4 by immunohistochemistry staining (Figure 7C). In accordance with the result from HPA database, the expression levels of IRS4 in ovarian tumor samples were significantly elevated (P = 0.0029) compared to normal control samples (Figure 7D).

To further assess the relationship between IRS4 expression and tumor progression, we analyzed clinical data from over 600 ovarian cancer patients (http://www.kmplot.com) and plotted the overall survival curves for both the IRS4-high and the IRS4-low cohorts. The result demonstrated that a lower expression of IRS4 was correlated to longer overall survival in patients with ovarian cancer (Figure 7E). In conclusion, IRS4 was significantly overexpressed in ovarian cancers and its upregulation was inversely correlated with survival and prognosis in ovarian cancer patients.

## Discussion

Insulin receptor substrates (IRSs) are cytoplasmic adaptor proteins that participate in the signal transduction process of various receptor tyrosine kinases (RTKs). They link RTKs such as insulin receptor (IR) and insulin-like growth factor 1 receptor (IGF-IR) to downstream signaling molecules containing Src homology 2 (SH2) domains (34). Their C-terminus possesses multiple tyrosine residues, the phosphorylation of which can create docking sites and recruit downstream effector proteins containing SH2 domains, activating the diverse signaling cascade and regulating cellular processes such as proliferation, differentiation, survival, metabolism and insulin resistance.

The IRS family consists of four closely related members IRS1-IRS4 and two distant relatives IRS5/DOK4 and IRS6/DOK5. Although the members of the IRSs family are similar in overall structure and possess high homology, there are differences in tissue distribution, expression, subcellular distribution, and interactions with protein molecules containing the SH2 domain, which enable the six IRS proteins to have different biological characteristics and mediate different signal transduction pathways.

Combining mass spectrometry analysis with biochemical and biological approaches, we have revealed one of the IRS family members, IRS4, as a novel substrate of non-receptor tyrosine kinase FER. FER binds directly to IRS4 (Fig 2) and phosphorylates several tyrosine residues on IRS4 (Fig 3). The proximity labeling PUP-IT assay further confirmed that FER-mediated phosphorylation of Tyr779 on IRS4 is critical in recruiting PIK3R2/p85β, the key regulatory subunit of PI-3K kinase (Fig 5). While ectopic over-expression of FER in HEK293FT cells dramatically increased tyrosine phosphorylation and PIK3R2 recruitment of IRS4, CRISPR-Cas9 directed knockout or pharmacological inhibition of the endogenous kinase in ovarian cancer cells remarkably reduced tyrosine phosphorylation and PIK3R2 recruitment of IRS4. The current working model is present in Fig 6E.

IRS4, originally known as PY160, was first cloned and identified as a new member of the IRS family in the HEK293 cell line (35). It has a typical molecular structure for the IRS family, that is, a N-terminal pleckstrin homology (PH) domain, a phosphotyrosine binding (PTB) domain and a C-terminal domain. The PH domain is composed of 122 amino acids (residues 78-199), which are very important for protein-protein and protein-phospholipid interactions. It predominantly binds to membrane phospholipids or the acidic motifs of various proteins on the cell membrane, immobilizing IRS4 on the cell membrane and increasing its proximity to the transmembrane receptor (36). The PTB domain consists of 104 amino acids (residues 231-334) adjacent to the PH domain and primarily recognizes the asparagine-proline-glutamate-tyrosine phosphorylation (NPEpY) sequence located in near-membrane region of the insulin receptor β subunit (35, 37). By using a series of truncation mutants of IRS4, we have illustrated that N-terminal region of IRS4 (1-334) is key for IRS4’s association with kinase FER (Fig 2D). Although there is no NPEpY motif in the C-terminus of FER, the region which is involved in FER-IRS4 interaction, we do see a similar PGEpY motif. The FES and FER proteins distinguish themselves from other tyrosine kinases through an N-terminal FES/FER/CIP4 homology/Bin1/Amphiphysin/RVS (F-BAR) lipid-binding domain (5, 6), which has functions in regulating vesicular trafficking and endocytosis. Interestingly, although the capacity of FER to deform protein-free liposomes into tubules is relatively weak (38), the capacity of its F-BAR domain and the F-BAR extension domain (FX) to strongly and specifically bind phosphatidic acid (PA) in the plasma membrane has been well characterized (39). A recent study from O’Connor’s team has reported co-localization and co-immunoprecipitation between FER and IGF-1R, the upstream RTK of IRS4 (24). In fact, IGF-1R was ranked 22^nd^ in our MS dataset (Fig 1C). It will therefore be very enlightening to further investigate the molecular interplay among IGF-1R, FER, and IRS4, and their impact on activating their downstream signaling pathways.

IRS1 and IRS2 possess specific tyrosine residues in their C-terminal tails, the phosphorylation of which facilitates the recruitment of SH2-containing tyrosine phosphatase SHP2. This provides negative feedback regulation via dephosphorylating tyrosine residues on the C-terminal tails of IRS1 and IRS2, which are responsible for downstream effectors recruitment and signaling propagation (40). Unlike IRS1 and IRS2, IRS4 has no SHP2 binding motif and does not bind to SHP2 (41). This unique feature allows IRS4 to maintain constitutive hyperactivation of the PI3K-AKT signaling pathway (41–43), leading to growth factor-independent cell proliferation and tumorigenesis in mammary epithelial cells (44). This was also the case in the ovarian cancer cell line, OVCAR-5, with an aberrantly high expression of IRS4 (Fig 4A). Genetic ablation of IRS4 with CRISPR-Cas9 almost completely abolished the activation of AKT kinase (Fig 4B) and dramatically delayed cell proliferation (Fig 4C). Interestingly, these cells were very sensitive to inhibitors against the PI3K-AKT-mTOR pathway (Fig 4F-H). This suggests that ovarian cells with high IRS4 expression may depend on IRS4-mediated PI3K-AKT activation for proliferation and survival. Any pharmacological perturbation of this pathway would benefit therapeutic outcomes for patients suffered from this deadly disease.

There are 7 Y-X-X-M motifs on IRS4 (Tyr-487, −700, −717, −743, −779, −828, −921), which have been speculated as potential binding sites for the regulatory subunit of PI-3K. Our results from mass spectrometry and site-directed mutagenesis analysis revealed 5 Tyr residues as potential substrates for FER kinase: Tyr-656, −779, −828, −847 and −921 (Fig 2C-E). Proximity labeling PUP-IT assays further demonstrated that Tyr779 was the major site responsible for PIK3R2/p85β recruitment on IRS4 (Fig 5B-F) and this result was confirmed by co-immunoprecipitation (Fig 5H). Our data provides compelling evidence to decipher the molecular details on PIK3R2/p85β association with IRS4.

Besides IRS4 and IGF-1R, our mass spectrometry analysis captured several interesting FER-interacting hits with functions of solute transportation, including SLC25A5, SLC25A6 SLC3A2, ATP1A1 and ATP2A2 (Fig 1C). Notably, tyrosine phosphorylation has been reported as an important layer of regulation for transporter proteins’ stability (45) and activity (46–49). We also identified CTLC (Clathrin Heavy Chain) and COPA (COPI Coat Complex Subunit Alpha) as potential FER associated proteins, the functions of which are involved in both Clathrin-dependent and -independent intracellular trafficking (Fig 1C). In addition, regulators in cytosol-nucleus shuttle, for example XPO1 and IPO4, were also ranked high in the hit list (Fig 1C). Considering the well documented function of FER kinase in vesicle trafficking and cell motility, these findings will definitely shed new light on molecular mechanism on FER’s function.

In summation, our study has demonstrated IRS4 as a novel substrate for non-receptor tyrosine kinase FER. This kinase-substrate regulatory mode between FER and IRS4, which leads to PIK3R2 recruitment and AKT activation, is critical for ovarian tumor cell growth. This work expounds on the versatile functions of the FER kinase, especially within ovarian cancer and highlights the unmet need to develop a small molecule inhibitor of the kinase to benefit patients.

## Materials and Methods

### Cell culture and chemical reagents

Human embryonic kidney 293FT cells, ovarian carcinoma-derived cell lines (OVCAR-5, OVK-18, COLO-316, IGOV-1, OVCAR-3, PA-1, CAOV4, CAOV-3, CAOV1, A2780 and SK-OV-3), immortalized human ovarian surface epithelial cell lines (HOSE 11-12 and HOSE 6-3) were cultured in Dulbecco’s modified Eagle’s medium (DMEM, Cellgro) supplemented with 10% fetal bovine serum (FBS, Cellgro), 100 units/ml penicillin and 100 μg/ml streptomycin. Cells were maintained at 37°C in 5% CO2.

AKT inhibitor LY294002 (S1105), Anti-cancer Metabolism Compound Library (L5700) and TAE684 (S1108) were purchased from Selleck.

### Plasmids

Mammalian expression plasmids used in this study were as follows: pMSCV-FER (a gift from Prof. Peter A. Greer, Queen’s University, Canada), pEGFP-FER (a gift from Prof. Toshiki Itoh, Kobe University, Japan), FPC1-myc-IRS4 (IRS4 full length and IRS4 truncation mutants 1-400, 1-550, 1-699, 1-800, 1-1093 and 200-1257 were gifts from Prof. Kensaku Mizuno, Tohoku University, Japan), pEYFPC1-IRS4 (IRS4 truncation mutants 1-334, 335-400, 200-400 and 401-1257 were gifts from Prof. Kensaku Mizuno, Tohoku University, Japan), pUSE-SRC, pcDNA6-HCK, pHAGE-FYN, pcDNA3.1-LCK, pLPC-BRK, pWZL-BTK, PX330-IRS4-sgRNA-Cas9-GFP, PX330-FER-sgRNA-Cas9-GFP, PUP-IT (pTet3G-Bio-PupE-IRES-BFP and IRS4-PafA-IRES-puro-GFP). FER cDNA (NM_001308028) and IRS4 cDNA (NM_003604) have been used in this study.

### Cell transfection and infection

We followed manufacture protocol of Mirus (TransIT-2020, Mirus Bio) to perform transient transfection. Briefly, cells were plated in a 6-well plate 24 hours prior to transfection. When cells reach ~75% confluence, prepare Mirus:plasmid complexes in Opti-MEM I Reduced Serum Medium (Gibco) and add them into each well. 24 hours later, cells were harvested and lysed for immunoblotting or immunoprecipitation assays.

Cell line with gene stable expression was established by lentiviral infection, followed by GFP sorting or puromycin selection. In brief, lentivirus was generated in HEK293FT cells by co-transfecting gene-containing plasmids, deltaR8.91 and VSVG at a ratio of 3:2:1. 48 to 72 hours later, supernatants were collected and passed through 0.45 um filters to remove cell debris. Cleared virus was then added to cells cells to be infected in the presence of polybrene. Infected cells were either sorted by GFP or selected by puromycin. The effectiveness of infection was confirmed by flow cytometry or immunoblotting with according antibody.

### Protein expression and purification

GST-tagged FER (1-446) and GST-tagged FER (447-End) were expressed in Escherichia coli BL21 (DE3). The cells were cultured at 37°C until the OD reached 0.8-1.0 and were induced with 0.3 mM isopropyl β-d-thiogalactoside (IPTG) in LB medium at 16°C overnight. Bacteria were lysed in lysis buffer (50 mM Tris-HCl, pH 7.5, 250 mM NaCl, 1 mM DTT and 1× complete protease inhibitor) by high-pressure homogenizer. After centrifugation, the supernatant was incubated with GST beads at 4°C for 2 hours. After washing, GST-tagged proteins were eluted with 10 mM reduced glutathione. Protein concentration was measured using the Bradford assay. Protein purity was assessed by SDS-PAGE and coomassie blue staining.

### Immunoblotting and immunoprecipitation assay

Cells were lysed in lysis buffer (20 mM HEPES pH 7.5, 150 mM NaCl, 1% Nonidet P-40, 1 mM sodium orthovanadate and 1× complete protease inhibitor cocktail from Roche) at 4°C for 15 minutes. Total protein concentration was determined by Bradford assay.

For immunoblotting, cellular proteins were harvested, separated by SDS-PAGE and transferred onto nitrocellulose membranes. Membranes were blocked in 2.5% BSA in TBST (TBS/Tween 20: 20 mM Tris HCl, pH 7.6, 136 mM NaCl, and 0.1% Tween 20) for 1 hour at room temperature on a shaker and incubated with primary antibody at 4°C overnight. Proteins were detected with horseradish peroxidase (HRP)-conjugated secondary antibodies (Jackson Laboratory) and ECL (Pierce).

For immunoprecipitation, precleared cell extracts were incubated with the indicated antibody for 4 hours at 4°C with rotation followed by 1 h of pull-down by 1:1 protein A/G agarose beads. Immunoprecipitates were washed with lysis buffer three times before electrophoresis.

The primary antibodies used in this study were as followed: 4G10 (Millipore); Myc (9E10); FER, phospho-Erk1/2, total Erk1/2, phospho-Ser473 AKT and total AKT (Cell Signaling Technology); phospho-p38 (Promega); total p38 (Santa Cruz Biotechnology); pY402 FER (Millipore); IRS4 and β-actin (Sigma); GAPDH (Novus Biologicals). The beads used in this assay were as followed: Streptavidin Magnetic Beads (NEB), EZview(TM) Red anti-c-Myc affinity gel (Sigma), protein A sepharose and protein G sepharose (GE).

### Cell proliferation assay

Cell Titer-Glo (CTG) luminescent cell viability assay (Promega) was used to evaluate the role of IRS4 in ovarian cancer cell proliferation. In brief, 1.5 × 10^3^ OVCAR-5 cells per well were seeded in a 96-well plate and grown for indicated time intervals. CTG reagent was added to each well and mixed for ~15 min on an orbital shaker to induce cell lysis followed by luminescence reading.

### CRISPR-Cas9 system for gene knockout

To generate IRS4 and FER knockout ovarian cancer cell lines using CRISPR-Cas9 system, the CRISPR sgRNA database (https://www.genscript.com/gRNA-database.html?src=leftbar) was applied to generate sgRNAs for each gene. The selected sgRNAs were then subcloned into PX330-Cas9-GFP plasmid, followed by transient transfection into ovarian cancer cell lines and FACS sorting for GFP-positive single clone. The knockout effect was confirmed by western blotting analysis against relevant antibodies.

The sgRNAs used were: IRS4 sgRNA#1 seq (5’-CCATCGCGAAGTATTCGTCT-3’), IRS4 sgRNA#2 seq (5’-TATAGGGTGATCACGCGCCG-3’), IRS4 sgRNA#3 seq (5’-CCGGCTGTGTCTAACCGACG-3’), FER sgRNA#1 seq (5’-TCTATTCGTCATTCAATTGC-3’), FER sgRNA#4 seq (5’-AGAGTTTGATACTTCCTTAC-3’) and FER sgRNA#5 seq (5’-CCGACATTTGAATCTCTTTA-3’)

### PUP-IT (pupylation-based interaction tagging) assay

The experimental procedure was modified based on previous study (31). To generate inducible Pup (iPup) cell lines, we first produced lentivirus with Pup (E) plasmid Bio-Pup (E)-IRES-BFP within the Tet-On 3G inducible expression system (Clontech 631168), and infected OVCAR-5 cells for 48 hours. Subsequently, doxycycline (final concentration 2 μg/ml Selleck, S4136) was added into the culture medium for another 24 hours. BFP-positive cells were then sorted into 96-well plates by flow cytometry for single clone selection. It takes ~3 weeks for cell re-population. After adding doxycycline (final concentration 2 μg/ml) and biotin (final concentration 4 μM) for 24 hours, the BFP expression of each clone was confirmed by flow cytometry, and the expression and modification of Bio-Pup (E) in BFP-positive cells was also confirmed by western blotting.

In order to further stably express IRS4 (WT)-PafA or IRS4 (YF)-PafA in iPup OVCAR-5 cells, we subcloned IRS4-WT or IRS4-YF into the PafA-IRES-puro-EGFP plasmid, respectively, and produced lentivirus to infect iPup OVACR5 cells for 48hours. Cells were placed under puromycin selection (final concentration 2μg/ml) for generating iPup OVCAR-5 cell lines which stably express IRS4 (WT)-PafA or IRS4 (YF)-PafA, respectively. The expression of IRS4 (WT)-PafA and IRS4 (YF)-PafA was confirmed by western blotting analysis.

IRS4 (WT)-PafA or IRS4 (YF)-PafA expressed iPup OVCAR-5 cells were then grown in 10 cm dishes. We added doxycycline (final concentration 2 μg/ml) and biotin (final concentration 4 μM) to the medium in advance, and induce expression in cells for 24 hours. Then, we harvested cells, and followed the protocol in (31) to prepare sample for mass spectrometry analysis.

Particularly in this study, we compared the binding protein differences between IRS4 (WT) and IRS4 (YF) in OVCAR-5 cells. To obtain reliable and quantitative measurement, each group of samples was triplicated. To analyze the different proteins bound to IRS4 (WT) or IRS4 (YF) in OVCAR-5 cells, we calculated the fold change of LFQ intensity and used the T test to calculate the P-value. Fold change > 2.3 and P-value < 0.05 would be regarded as differences and statistical significance. We used GraphPad Prism to draw the relevant volcano maps.

### Sample preparation, digestion and mass spectrometry

For identification of phosphorylated tyrosine residues and interacting proteins by mass spectrometry, immunoprecipitates were prepared first as described above. Samples were subjected to SDS–PAGE gel, followed by in-gel trypsin digestion. In brief, gel bands were excised, washed and dehydrated with 100% acetonitrile. Proteins inside the gel were reduced, alkylated and finally digested with trypsin overnight at 37°C. The mixture of peptide fragments were extracted with 50% acetonitrile and 1% trifluoroacetic acid followed by 100% acetonitrile. Peptides were vacuum-dried and re-suspended for following mass spectrometry characterization. When samples were subjected to on-beads digestion, the beads in immunoprecipitation were digested with trypsin overnight at 37°C. After cleaning, peptides were vacuum-dried and re-suspended for following mass spectrometry characterization.

Mass spectrometry analysis was performed at the Proteomics Facility in Shanghaitech University. An Easy-nLC 1000 system coupled to a Q Exactive HF (both from Thermo Scientific) was used to separate and analyze peptides. The raw data were processed and searched with MaxQuant 1.5.4.1 with MS tolerance of 4.5 ppm, and MS/MS tolerance of 20 ppm. The UniProt human protein database (release 2016_07, 70630 sequences) and database for proteomics contaminants from MaxQuant were used for database search.

### Animal work

All study protocols involving mice were approved by the Institutional Animal Care and Use Committee of the ShanghaiTech University and conducted in accordance with governmental regulations of China for the care and use of animals. In the subcutaneous injection model, 1×10^6^ OVCAR-5 IRS4-KO cells with ectopic expression of EV (n=5), IRS4-WT (n=4) or IRS4-5YF (n=7), respectively, were suspended in 100 μL of 1:1 mixture with DMEM and growth factor-reduced Matrigel (BD Biosciences) and subcutaneously injected into NSG mice. Subcutaneous tumor growth was monitored periodically by measuring tumor volume (in cubic millimeters, formula: volume=width^2^×length/2) with calipers.

### Immunohistochemistry staining

Paraffin-embedded tissues were sectioned and stained with H&E or specific immunohistochemical stains. Stained slides were digitally scanned using the Aperio ScanScope software. H score was used for statistical analysis and calculated as positive staining percentage multiplied by staining strength (50). Both positive and negative control slides were included. The IRS4 primary antibody used in this assay was from Sigma.

### Statistics

Statistics were performed using a standard Student’s T-test or two way ANOVA multiple comparisons.

## Acknowledgements

This work was supported by the Ministry of Science and Technology of China (2018YFC1004603 to G.F.), the National Natural Science Foundation of China (31872831 to G.F.), Science and Technology Commission of Shanghai Municipality (19JC1413800 to G.F.), the Shanghai Pujiang program (18PJ1407900 to G.F.), the Shanghai Shuguang Program (19SG55 to G.F.) and ShanghaiTech University Startup grant (to G.F.). We thank Dr. Haopeng Wang (ShanghaiTech University) for helpful comments in the preparation of the manuscript.

## Contributions

GF designed the study. YZ, XX, JZ, YW, LC, SC, LH and XZ performed the experiments; JC and DL collected the human patient samples; QZ and PH analyzed MS data. YZ, MZ and GF interpreted the data; YZ and GF wrote the paper, with the help from XX.

The authors declare that they have no competing interests.

## Notes

### Competing Interest Statement

The authors have declared no competing interest.

